# Deep Learning for Quality Control of Subcortical Brain 3D Shape Models

**DOI:** 10.1101/402255

**Authors:** The ENIGMA Consortium, Dmitry Petrov, Boris A. Gutman, Egor Kuznetsov, Theo G.M. van Erp, Jessica A. Turner, Lianne Schmaal, Dick Veltman, Lei Wang, Kathryn Alpert, Dmitry Isaev, Artemis Zavaliangos-Petropulu, Christopher R.K. Ching, Vince Calhoun, David Glahn, Theodore D. Satterthwaite, Ole Andreas Andreassen, Stefan Borgwardt, Fleur Howells, Nynke Groenewold, Aristotle Voineskos, Joaquim Radua, Steven G. Potkin, Benedicto Crespo-Facorro, Diana Tordesillas-Gutirrez, Li Shen, Irina Lebedeva, Gianfranco Spalletta, Gary Donohoe, Peter Kochunov, Pedro G.P. Rosa, Anthony James, Udo Dannlowski, Bernhard T. Baune, Andr Aleman, Ian H. Gotlib, Henrik Walter, Martin Walter, Jair C. Soares, Stefan Ehrlich, Ruben C. Gur, N. Trung Doan, Ingrid Agartz, Lars T. Westlye, Fabienne Harrisberger, Anita Riecher-Rössler, Anne Uhlmann, Dan J. Stein, Erin W. Dickie, Edith Pomarol-Clotet, Paola Fuentes-Claramonte, Erick Jorge Canales-Rodrguez, Raymond Salvador, Alexander J. Huang, Roberto Roiz-Santiaez, Shan Cong, Alexander Tomyshev, Fabrizio Piras, Daniela Vecchio, Nerisa Banaj, Valentina Ciullo, Elliot Hong, Geraldo Busatto, Marcus V. Zanetti, Mauricio H. Serpa, Simon Cervenka, Sinead Kelly, Dominik Grotegerd, Matthew D. Sacchet, Ilya M. Veer, Meng Li, Mon-Ju Wu, Benson Irungu, Esther Walton, Paul M. Thompson, for the ENIGMA consortium

## Abstract

We present several deep learning models for assessing the morphometric fidelity of deep grey matter region models extracted from brain MRI. We test three different convolutional neural net architectures (VGGNet, ResNet and Inception) over 2D maps of geometric features. Further, we present a novel geometry feature augmentation technique based on parametric spherical mapping. Finally, we present an approach for model decision visualization, allowing human raters to see the areas of subcortical shapes most likely to be deemed of failing quality by the machine. Our training data is comprised of 5200 subjects from the ENIGMA Schizophrenia MRI cohorts, and our test dataset contains 1500 subjects from the ENIGMA Major Depressive Disorder cohorts. Our final models reduce human rater time by 46-70%. ResNet outperforms VGGNet and Inception for all of our predictive tasks.

## 1 Introduction

Quality control (QC) has become one of the main practical bottlenecks in bigdata neuroimaging. Reducing human rater time via predictive modeling and automated quality control is bound to play an increasingly important role in maintaining and hastening the pace of scientific discovery in this field. Recently, the UK Biobank publicly released over 10,000 brain MRIs (and planning to release 90,000 more); as other biobanking initiatives scale up and follow suit, automated QC becomes crucial.

In this paper, we investigate the viability of deep convolutional neural nets for automatically labeling deep brain regional geometry models of failing quality after their extraction from brain MR images. We compare the performance of VGGNet, ResNet and Inception architectures, investigate the robustness of probability thresholds, and visualize decisions made by the trained neural nets. Our data consists of neuroimaging cohorts from the ENIGMA Schizophrenia and Major Depressive Disorder working groups participating in the ENIGMA-Shape project [1]. Using ENIGMAs shape analysis protocol and rater-labeled shapes, we train a discriminative model to separate FAIL(F) and PASS(P) cases. Features are derived from standard vertex-wise measures.

For all seven deep brain structures considered, we are able to reduce human rater time by 46 to 70 percent in out-of-sample validation, while maintaining FAIL recall rates similar to human inter-rater reliability. Our models generalize across datasets and disease samples. Our models’ decision visualization, particularly ResNet, appears to capture structural abnormalities of the poor quality data that correspond to human raters’ intuition.

With this paper, we also release to the community the feature generation code based on FreeSurfer outputs, as well as pre-trained models and code for model decision visualization.

## 2 Methods

Our goal in using deep learning (DL) for automated QC differs somewhat from most predictive modeling problems. Typical two-class discriminative solutions seek to balance misclassification rates of each class. In the case of QC, we focus primarily on correctly identifying FAIL cases, by far the smaller of the two classes **(Table 1)**. In this first effort to automate shape QC, we do not attempt to eliminate human involvement, but simply to reduce it by focusing human rater time on a smaller subsample of the data containing nearly all the failing cases.

**Table 1:**
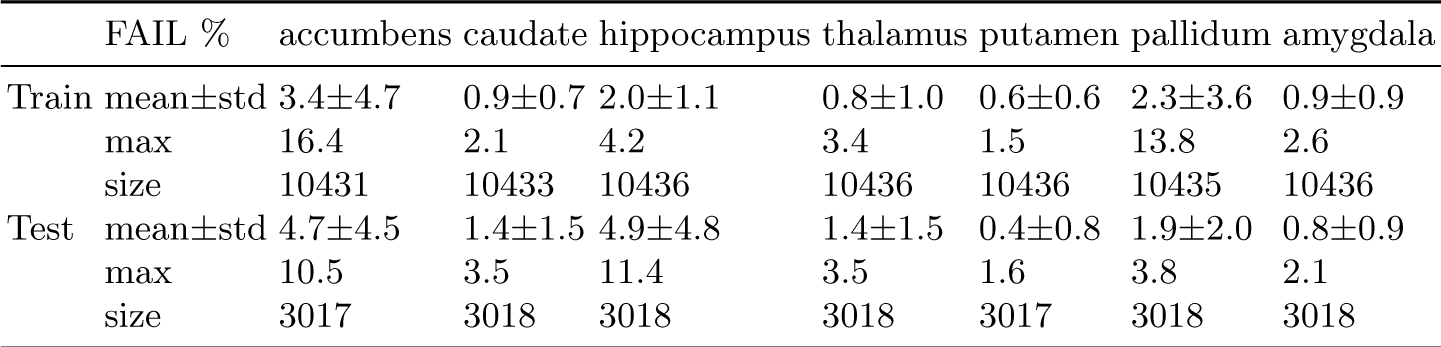
Overview of FAIL percentage mean, standard deviation and maximum for each site. Minimum is equal to 0 for all regions and sites except for hippocampus on train (FAIL percentage 5%). Sample sizes for each ROI vary slightly due to FreeSurfer segmentation failure.

### 2.1. MRI processing and shape features

Our deep brain structure shape measures are computed using a previously described pipeline [2] [3], available via the ENIGMA Shape package. Briefly, structural MR images are parcellated into cortical and subcortical regions using FreeSurfer. Among the 19 cohorts participating in this study, FreeSurfer versions 5.1 and 5.3 were used. The binary region of interest (ROI) images are then surfaced with triangle meshes and spherically registered to a common regionspecific template [4]. This leads to a one-to-one surface correspondence across the dataset at roughly 2,500 vertices per ROI. Our ROIs include the left and right thalamus, caudate, putamen, pallidum, hippocampus, amygdala, and nucleus accumbens. Each vertex *p* of mesh model is *ℳ* endowed with two shape descriptors:

Medial Thickness, *D*(*p*) = ‖ *c*_*p*_*-p* ‖, where *c*_*p*_ is the point on the medial curve *c* closest to *p*.

*LogJac*(*p*), Log of the Jacobian determinant *J* arising from the template mapping, *J*: *T*_*φ*(*p*)_ *ℳ* _*t*_ *→ T*_*p*_. *ℳ*

Since the ENIGMA surface atlas is in symmetric correspondence, i.e., the left and right shapes are vertex-wise symmetrically registered, we can combine the two hemispheres for each region for the purposes of predictive modeling. Though we assume no hemispheric bias in QC failure, we effectively double our sample.

The vertex-wise features above are augmented with their volume-normalized counterparts: 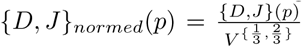 Given discrete area elements of the template at vertex *p, A*_*t*_(*p*), we estimate volume as 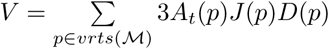 We normalize our features subject-wise by this volume estimate to control for subcortical structure size.

### 2.2. Human quality rating

Human-rated quality control of shape models is performed following the ENIGMAShape QC protocol (enigma.usc.edu/ongoing/enigma-shape-analysis). Briefly, raters are provided with several snapshots of each region model as well as its placement in several anatomical MR slices. A guide with examples of FAIL (QC=1) and PASS (QC=3) cases is provided to raters, with an additional category of MODERATE PASS (QC=2) suggested for inexperienced raters. With sufficient experience, the rater typically switches to the binary FAIL/PASS rating. In this work, all QC=2 cases are treated as PASS cases, consistent with ENIGMA shape studies.

### 2.3. Feature mapping to 2D images

Because our data resides on irregular mesh vertices, we first interpolate the features from an irregular spherical mesh onto an equiangular grid. The interpolated feature maps are then treated as regular 2D images by Mercator projection. Our map is based on the medial curve-based global orientation function (see [5]), which defines the latitude (*θ*) coordinate, as well as a rotational standardization of the thickness profile *D*(*p*) to normalize the longitudinal (*φ*) coordinate. The resulting map normalizes the 2D appearance of *D*(*p*), setting the poles to lie at the ends of the medial curve. In practice, the re-sampling is realized as matrix multiplication based on trilinear mesh interpolation, resulting in a 128 *×* 128 image for each measure.

### 2.4. Data augmentation

Although our raw sample of roughly 13,500 examples is exceptionally large by the standards of neuroimaging, this dataset may not be large enough to train generalizable CNNs. Standard image augmentation techniques, e.g. cropping and rotations, are inapplicable to our data. To augment our sample of spherically mapped shape features, we sample from a distribution of spherical deformations, i.e. changes in the spherical coordinates of the thickness and Jacobian features. To do this, we first sample from a uniform distribution of vector spherical harmonic coefficients ***B***_*lm*_, ***C***_*lm*_, and apply a heat kernel operator [4] to the generated field on *T* 𝒮 ^2^. Change in spherical coordinates is then defined based on the tangential projection of the vector field, as in [4]. The width *σ* of the heat kernel defines the level of smoothness of the resulting deformation, and the maximum point norm *M* defines the magnitude. In practice, each random sampling is a composition of a large magnitude, smooth deformation (*σ* = 10^−1^, *M* = 3*×*10^−1^) and a smaller noisier deformation (*σ* = 10^−2^, *M* = 3*×*10^−2^). Once the deformation is generated, it is applied to the spherical coordinates of the irregular mesh, and a new sampling matrix is generated, as above.

### 2.5. Deep learning models

We train VGGNet [6], ResNet [7] and Inception [8] architectures on our data. We chose these architectures as they perform well in traditional image classification problems and are well-studied.

### 2.6. Model decision visualization

Deep learning models tend to learn superficial statistical patterns rather than high-level global or abstract concepts [9]. As we plan to provide a tool that both (1) classifies morphometric shapes, and (2) allows a user to visualize what the machine perceives as a ‘FAIL’, model decision visualization is an important part of our work. Here, we use Prediction Difference Analysis [10], and Grad-CAM [11] to visualize ‘bad’ and ‘good’ areas for each particular shape in question.

### 2.7. Predictive model assessment

We use two sets of measures to evaluate the performance of our models. To assess the validity of the models’ estimated ‘FAIL’ probabilities, we calculate the area under the ROC curve (ROC AUC). We also use two supplementary measures: FAIL-recall and FAIL-share. In describing them below, we use the following definitions. TF stands for TRUE FAIL, FF stands for FALSE FAIL, TP stands for TRUE PASS, and FP stands for FALSE PASS. Our first measure, 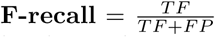, shows the proportion of FAILS that are correctly labeled by the predictive model with given probability threshold. The second measure, 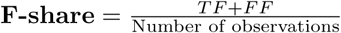, shows the proportion of the test sample labeled as FAIL by the model. Ideal models produce minimal **F-share**, and an **F-recall** of 1 for a given set of parameters.

## 3 Experiments

For each of the seven ROIs, we performed three experiments defined by three DL models (VGGNet-, ResNet- and Inception-like architecture).

### 3.1. Datasets

Our experimental data from the ENIGMA working groups is described in **Table 1**. Our predictive models were trained using 15 cohorts totaling 5218 subjects’ subcortical shape models from the ENIGMA-Schizophrenia working group. For a complete overview of ENIGMA-SCZ projects and cohort details, see [12].

To test our final models, we used data from 4 cohorts in the Major Depressive disorder working group (ENIGMA-MDD), totaling 1509 subjects, for final out-of-fold testing. A detailed description of the ENIGMA-MDD sites and its research objectives may be found here [13].

### 3.2. Model validation

All experiments were performed separately for each ROI. The training dataset was split into two parts referred to as ‘TRAIN GRID’ (90% of train data) and ‘TRAIN EVAL.’ (10% of the data). The two parts contained data from each ENIGMA-SCZ cohort, stratified by the cohort-specific portion of FAIL cases.

Each model was trained on ‘TRAIN GRID’ using the original sampling matrix and 30 augmentation matrices resulting in 31x augmented train dataset. We also generated 31 instances of each mesh validation set using each sampling matrix and validated models’ ROC AUC on this big validation set during the training.

As models produce probability estimates of FAIL (*P*_*FAIL*_), we studied the robustness of the probability thresholds for each model. To do so, we selected *P*_*FAIL*_ values corresponding to regularly spaced percentiles values of **F-share**, from 0.1 to 0.9 in 0.1 increments. For each such value, we examined **F-recall** the evaluation set.

Final thresholds were selected based on the lowest **F-share** on the TRAIN EVAL set, requiring that **F-recall** *≥* 0.8, a minimal estimate of inter-rater reliability. It is important to stress that while we used sample distribution information in selecting a threshold, the final out-of-sample prediction is made on an individual basis for each mesh.

## 4 Results

Trained models were deliberately set to use a loose threshold for FAIL detection, predicting 0.2-0.5 of observations as FAILs in the ‘TRAIN EVAL’ sample. These predicted FAIL observations contained 0.85-0.9 of all true FAILs, promising to reduce the human rater QC time by 50-80%. These results largely generalized to the test samples: **Table 2** shows our best model and the threshold performance for each ROI. When applied to the test dataset, the models indicated modest over-fitting, with the amount of human effort reduced by 46-70%, while capturing 76-94% of poor quality meshes. The inverse relationship between FAIL percentage and F-share (**Figure 1**) may indicate model failure to learn generalizable features on smaller number of FAIL examples. ROC AUC and F-recall performance generalize across the test sites. Since 68% of our test dataset is comprised of the Munster cohort, it is important that overall test performance is not skewed by it.

**Table 2:** Test performance of the best models for each region. ResNet performs the best in all cases. Overall models’ performance generalizes to out-of-sample test data.

| ROI | Model | Eval<br>AUC | Test<br>AUC | Eval<br>F-share | Test<br>F-share | Eval<br>F-recall | Test<br>F-recall |
| --- | --- | --- | --- | --- | --- | --- | --- |
| Accumbens | ResNet | 0.86 | 0.8 | 0.3 | 0.35 | 0.83 | 0.78 |
| Amygdala | ResNet | 0.8 | 0.75 | 0.5 | 0.54 | 0.92 | 0.8 |
| Caudate | ResNet | 0.9 | 0.84 | 0.2 | 0.3 | 0.82 | 0.78 |
| Hippocampus | ResNet | 0.85 | 0.93 | 0.3 | 0.36 | 0.81 | 0.92 |
| Pallidum | ResNet | 0.86 | 0.91 | 0.3 | 0.32 | 0.81 | 0.91 |
| Putamen | ResNet | 0.88 | 0.7 | 0.3 | 0.52 | 0.93 | 0.76 |
| Thalamus | ResNet | 0.8 | 0.87 | 0.4 | 0.47 | 0.82 | 0.94 |

**Fig. 1:**
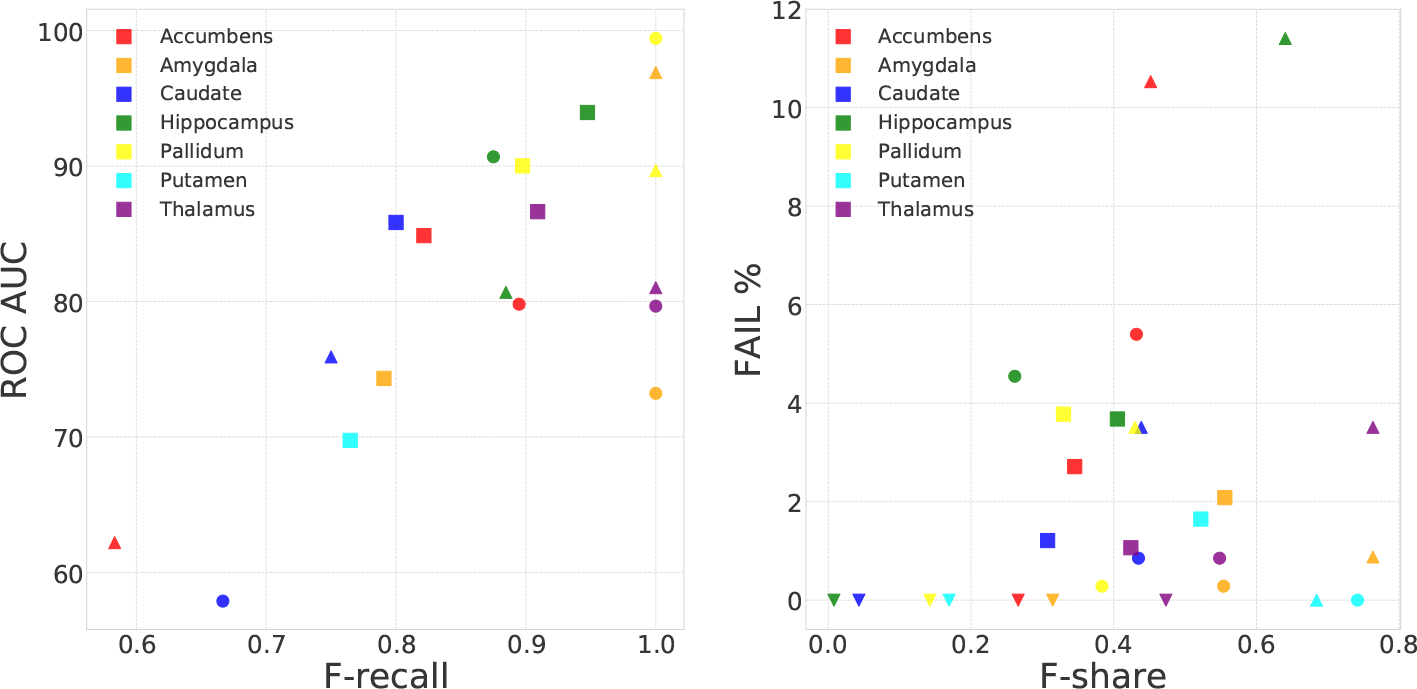
Scatter plots of F-recall vs. ROC AUC on test datasets and F-share vs. proportion of predicted FAIL cases on test datasets (F-share). **Left:** F-recall vs ROC AUC. **Right:** Fail F-share vs FAIL percentage. F-share was calculated based on thresholds from **Table 2**. Mark size shows the dataset size. Mark shape represents dataset (site): ◯-CODE-Berlin (N=176); □ - Munster(N=1033); △-Stanford (N=105); ▽ - Houston(N=195)

Our experiments with decision visualization (see **Fig. 2**) indicate that in most FAIL cases, the attention heat map generated by Grad-CAM corresponds to human raters’ intuition while Prediction Difference Analysis tend to concentrate on local ‘bumps’ on shapes.

**Fig. 2:**
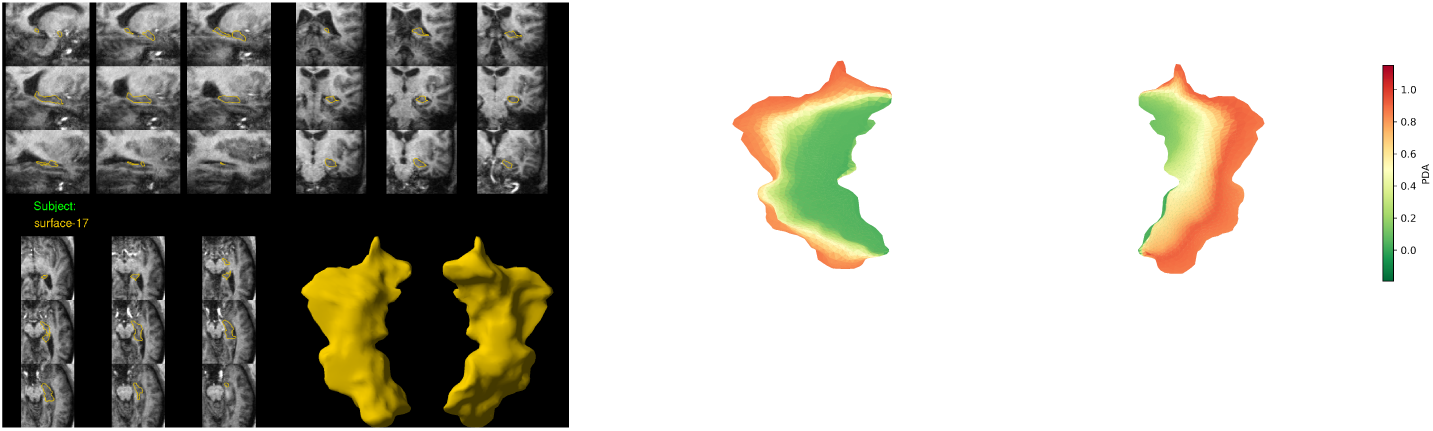
QC report for human raters (**left**) and decision visualization example based on Grad-CAM for the ResNet model (**right**). Red colors correspond to points maximizing the model’s FAIL decision in the last layer. Decision visualization corresponds to the observable deviations from underlying anatomical boundaries indicative of a “FAIL” rating according to an experienced rater.

## 5 Discussion and Conclusion

We have presented potential deep learning solutions for semi-automated quality control of deep brain structure shape data. We believe this is the first DL approach for detecting end-of-the-pipeline feature failures in deep brain structure geometry. We showed that DL can robustly reduce human visual QC time by 46-70% for large-scale analyses, for all seven regions in question, across diverse MRI datasets and populations. Qualitative analysis of models decisions shows promise as a potential training and heuristic validation tool for human raters.

There are several limitations of our work. Our planar projection of vertexwise features introduces space-varying distortions and boundary effects that can affect training, performance and visualization. Recently proposed spherical convolutional neural nets [14] may be useful to fix this issue. Second, our models’ decision visualization only partly matches with human raters’ intuition. In some cases, our models do not consider primary “failure” areas, as assessed by a human rater. Finally, our models are trained on purely geometrical features and do not include information on shape boundaries inside the brain. In rare cases, human raters pass shapes with atypical geometry because their boundaries look reasonable, and conversely mark normal-appearing geometry as failing due to poor a fit with the MR image. Incorporating intensity as well as geometry features will be the focus of our future work.

